# Discovery and binding site identification of a single-stranded DNA-binding protein that interacts with double-stranded DNA

**DOI:** 10.1101/2024.05.23.595439

**Authors:** Wang Yao, Tang Yajie, Jiang Zhihui, Ma Lei, Liu Yiwen, Lin Sen, Xu Yaoyu, Zhang Wenhui, Weng Shaoting

## Abstract

Scientists have conducted in-depth studies on single-stranded DNA (ssDNA) binding and protein interaction network of single-stranded DNA binding proteins (SSBs). SSBs are widely present in cells from lower prokaryotes to higher mammals and are important proteins for ssDNA protection. In-depth studies in recent years have revealed a wide range of interactions between SSBs and other proteins. Studies have shown that in addition to playing the role of ssDNA protector, SSBs are also important members of biochemical pathways such as DNA replication and recombination. In this study, a single-stranded DNA binding protein SSF encoded by *E. coli* F plasmid with highly conserved N-terminal region was demonstrated to have ssDNA binding ability similar to *E. coli* SSB through gel migration experiments. At the same time, the dsDNA binding ability of SSF was also demonstrated, and the dissociation constant of this interaction (in the range of microliters) was determined by Micro-scale Thermophoresis (MST). Through the above experiments, a previously unreported SSB with dsDNA binding ability was revealed, and the region interacting with dsDNA was determined to be the C-terminal unstructured region. This study lays a foundation for further understanding of the mechanism of bacterial conjugation conducted by F plasmid, as well as the potential dsDNA binding ability and biological significance of other SSBs.

## Introduction

Escherichia coli (*E.coli*) single stranded DNA binding protein (SSB) is a homologous tetramer with each subunit containing 178 amino acids and a molar mass of 18843.8 ^[1,2]^. And each subunit contains a highly conserved N-terminal DNA binding domain (also known as OB fold; residue 1-115) and an unstructured and poorly conserved C-terminal sequence (residues 116-177). SSB can bind to single stranded DNA (ssDNA) produced during DNA replication, repair, and recombination processes, and play a protective role for ssDNA. However, in recent years, an increasing number of studies have revealed that SSB is not only a guardian of ssDNA, but its highly conserved acidic C-terminal eight amino acid residues (SSB Ct) can also bind to over 12 different proteins ^[3]^. Kozlov *et al*. (2010) demonstrated through their research that the absence of eight residues in SSB-Ct can enhance the interaction between SSB and ssDNA ^[4]^.

Chase *et al*. (1983) discovered another ssDNA binding protein encoded by the *ssf* gene on the F plasmid in F+ *E.coli*, named SSF ^[5]^. Although there has been a lack of thorough research since its discovery, the alignment of its full-length sequence (179 amino acids) with the SSB sequence shows a high degree of conservatism in the OB domain^[6]^, but the similarity in the C-terminus region is minimal except for the eight amino acids at the end (**Figure 1**). It is interesting that the number of acidic and alkaline residues distributed in the C-terminal region of SSF are several times more than those of SSB (**Figure 1**), but there are currently no reports on the function and structure of the C-terminal region of SSF. Many aromatic SSB residues located in the N-terminal OB folding region, including Trp-40, Trp-54, Trp-88, His-55, Phe-60, and Tyr-70, previously identified as involved in ssDNA binding, are conserved in SSF ^[7]^. The residues involved in both monomer-monomer and dimer-dimer interactions in SSB also conserved in SSF ^[6,7]^. SSF can also support the survival of *E.coli* that is completely lacking the *ssb* gene ^[8]^, indicating that SSF is functionally similar to SSB.

**Figure 1:**
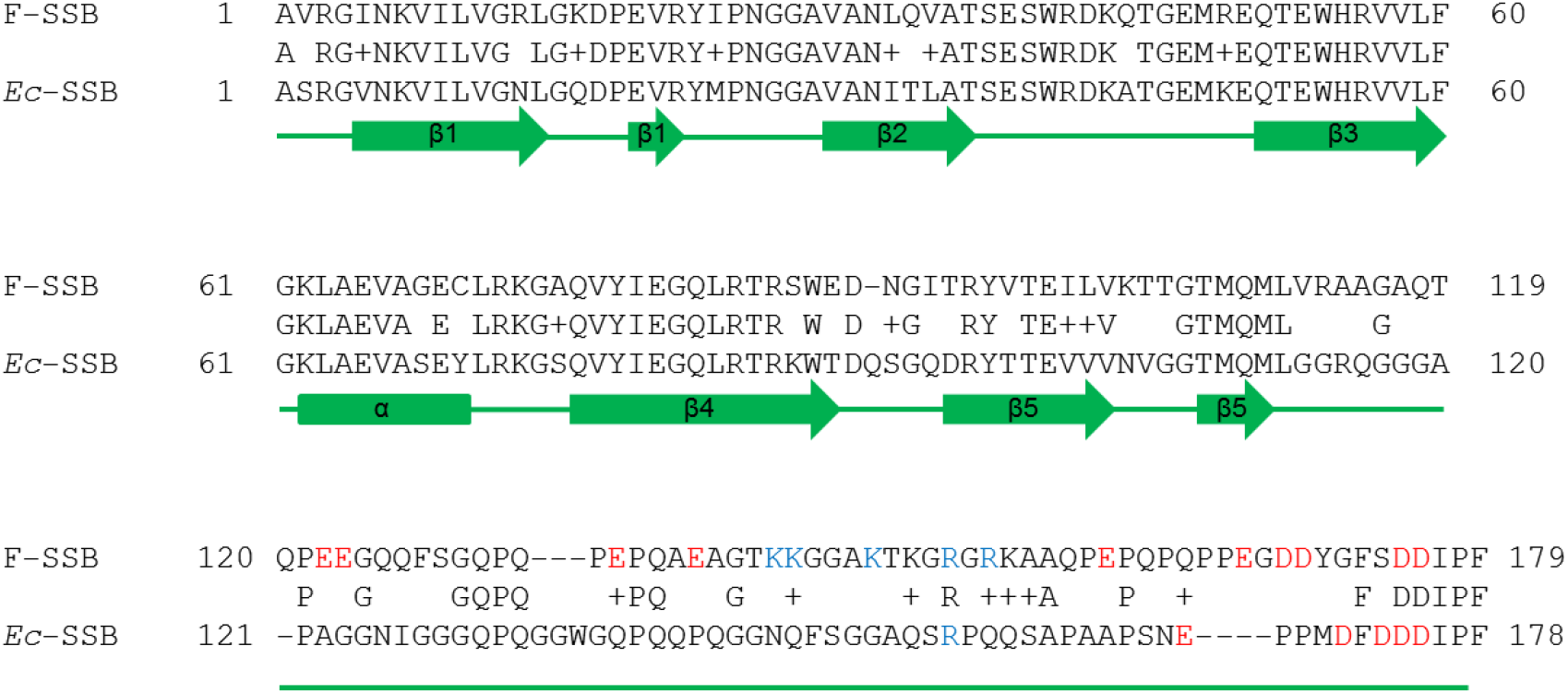
Comparison of full-length SSF and SSB sequences. The secondary structure of the protein and the acidic (red) and alkaline (blue) amino acids at the C-terminus were labeled.

This study reports for the first time the interaction properties between a single stranded DNA binding protein SSF and double stranded DNA (dsDNA) through biochemical experiments, and determines that the interaction region is at the C-terminus of SSF. SSF has the ability to bind both ssDNA and dsDNA, and after overexpression in cells, the complete dsDNA plasmid “disappear” in agarose gel electrophoresis. This study has opened up a new direction for SSB research.

## 1 Materials and Methods

### 1.1 Construction of expression vector and protein expression

pET11a plasmids containing wild-type ssf (GenBank: K00066.1), ssb (Genbank: J01704.1), Ubi-SSFc (CATCATCATCATCATCATATGCAGATCTTCGTGAAGACCCTGTGCGGTAAAACCATTACCCTGGAAGTGGAACCGTGCGACACCATCGAGAACGTTAAGGCGAAAATCCAAGATAAGGAAGGTATTCCGCCGTGCCAGCAACGTCTGATTTTCGCGTGCAAACAGCTGGAGGACGGCCGTACCCTGAGCGATTACAACATCCAAAAGTGCAGCACCCTGCACCTGGTTCTGGCGCTGGCGGGTGGCGGTGGCAGCGGTGGCAGCTTTAACGCGCGTCGTAAGCTGAAAGGCGCGATTCTGACCACCATGCTGGCGACCGCAGGTGCTCAGACTCAGCCGGAAGAGGGGCAACAGTTCAGCGGTCAGCCTCAGCCGGAACCACAGGCGGAGGCCGGTACGAAAAAAGGTGGCGCAAAAACGAAAGGCCGTGGACGTAAGGCCGCGCAGCCGGAGCCTCAGCCGCAACCGCCGGAGGGTGACGATTACGGGTTTTCAGACGATATCCCGTTC) and His6-SSFc (CATCATCATCATCATCATGCAGGTGCTCAGACTCAGCCGGAAGAGGGGCAACAGTTCAGCGGTCAGCCTCAGCCGGAACCACAGGCGGAGGCCGGTACGAAAAAAGGTGGCGCAAAAACGAAAGGCCGTGGACGTAAGGCCGCGCAGCCGGAGCCTCAGCCGCAACCGCCGGAGGGTGACGATTACGGGTTTTCAGACGATATCC CGTTC) are provided by GZL Bioscience Co. Ltd. All genes are inserted between the *Nde*I and *Bam*HI cleavage sites. Proteins are overexpressed by transforming the plasmids into BL21(λDE3)recA cells or using a cell-free protein expression kit (Cat. No: CF-EC-1000D; provided by GZL Bioscience Co. Ltd.). The procedures for cell-free protein expression are the same as previous reports ^[9,10]^. When expressing proteins *in vivo*, the bacteria are incubated with 1L LB liquid culture medium in 37℃ with agitation. 1 mM IPTG were added when OD_600_ reaches approximately 0.7, and then continue to incubate for 3 hours. Bacteria were collected by centrifugation and lysed for purification.

### 1.2 protein purification

Purification procedures of SSB were the same as reported by Mason *et al*. (2013) ^[11]^. Similar method was used to purify SSF: first, mix *E.coli* cells containing overexpressed SSF protein in a ratio of 0.07 g/mL with lysis buffer (50 mM Tris HCl, pH 8.0, 1 mM EDTA, 2 mM DTT, 0.5 mM PMSF), and use a high-pressure homogenizer to lyse cells at 12000 psi pressure. After centrifugation in 35000×g for 40 minutes, the supernatant containing SSF was mixed with 0.18 g/mL ammonium sulfate at 4℃. Stir for 30 minutes to precipitate SSF, and collect SSF by centrifugation at 35000×g for 30 minutes. Resuspend the precipitate in dialysis buffer (50 mM Tris HCl, pH 8.0, 1 mM EDTA, 1 mM DTT, 20 mM NaCl) and dialyse against 2 L of the same dialysis buffer in 4°C, then centrifuge in 35000×g for 30 minutes. Load the supernatant into a P11 cellulose column that was equilibrated with the same dialysis buffer. Wash the column material with 150 mL of the same dialysis buffer and elute target protein with a linear gradient of 0-1.4 M NaCl (360 mL). The purification of Ubi-SSFc and His6-SSFc was performed using HisTrap resin and AKTA protein purification system (Ge Healthcare Life Sciences). Firstly, wash the resin with washing buffer (20 mM sodium phosphate, 500 mM NaCl, pH 7.6), and then elute the protein with a linear imidazole gradient until a concentration of 500 mM is reached. Collect the sample with the highest purity. SSFΔ C62 (an SSF truncation protein that lacks the entire C-terminal sequence of the SSF and only contains the N-terminal OB region) was provided by GZL Bioscience Co. Ltd.

### 1.3 agarose gel shift assay

_SS_DNA and dsDNA binding ability of SSB and SSF were detected by agarose (1%) gel electrophoresis with 1 × TAE buffer. Mix dsDNA or ssDNA with certain concentrations (specific conditions are shown in the text) of SSB, SSF, Ubi-SSFc, and His6-SSFc in 10 mM Tris HCl (pH 7.6), 10 mM MgCl2, 1 mM DTT, and 40 mM NaCl, respectively. The sample was loaded into the agarose gel, and electrophoresis was carried out at 30 V for 3 hours in room temperature. After staining with SYBR Green (Bio Rad Co. Ltd), the DNA bands were observed. In order to investigate the ability of SSF to bind ssDNA and dsDNA synergistically, dT35 single stranded DNA was added in some experiments, and specific conditions are shown in the main text.

### 1.4 Microscale Thermophoresis

The Monolith NT 115 (Nanotemper Technologies Co. Ltd) instrument was used for the MST (Microscale Thermohoresis) experiment, and the data was collected and processed using NT analysis software. Mix 200 nM of full-length SSF and FITC (fluorescence isothiocyanate) labeled (Thermo Scientific Co. Ltd) His6-SSFc with 0.39 µM, 0.78 µM, 1.56 µM, 6.25 µM, 12.50 µM, 25.00 µM, 100.00 µM and 200.00 µM dsDNA hairpin (DNA sequence: GGC AAA TTA CTC CTG TAA TTT GCC) in PBS (pH 7.2) buffer, and then load the mixture into Monolith NT capillary (hydrophilic). Use blue LED light (80% power) to excite the fluorescence. Repeat the above experiment using full-length SSB and hairpin dsDNA for control.

### 1.5 *in vivo* dsDNA interaction assay

The pET plasmid containing the full-length *ssf* gene was transformed into the *E.coli* BL21(λDE3)recA. The culture was incubated at 37 ℃until OD_600_ reached 0.6, at which point 0.5 mM of IPTG was added for induction of SSF. The cells before and after induction were collected for small-scale plasmid extraction, and the effects of SSF on double stranded DNA plasmids were observed in agarose (1%) gel electrophoresis in 1× TAE buffer. Repeat the above experiment to observe the effects of His6-SSFc and full-length SSB on dsDNA plasmids.

## 2 Results

### 2.1 Expression and purification of full-length SSB, SSF和truncated SSF

To investigate the interaction between ssDNA, dsDNA and SSF, unlabeled full-length SSF was purified using ion exchange resin (**Figure 2A**). Similar purification procedures were used to purify the full-length SSB (**Figure 2A**). The N-terminal amino acid sequences of the two proteins mentioned above are highly similar, while the C-terminal sequence (SSFc) is almost completely different (**Figure 1**). The C-terminal sequence of SSB has been confirmed to have no interaction with DNA^[11-20]^. To explore the possibility of interaction between SSFc and DNA, we constructed Ubi-SSFc (SSFc fused with the C-terminus of a His6-tagged ubiquitin protein) and His6 labeled SSFc (His6-SSFc), and purified them using His-Trap resin (**Figure 2B** and **C**). The design of all protein constructs involved in this experiment are shown in **Figure 2D**.

**Figure 2:**
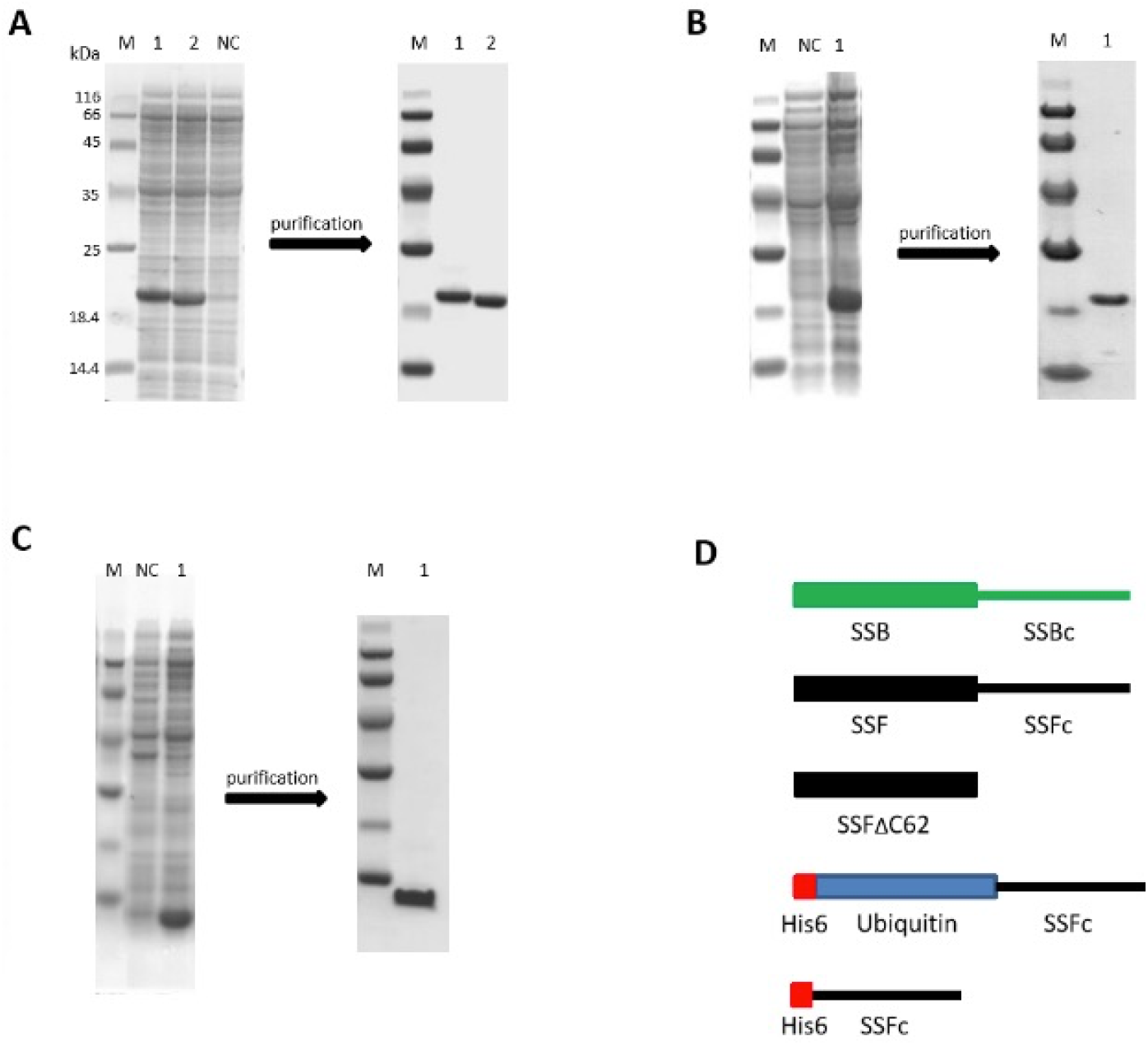
Expression and purification of full-length SSB, SSF, Ubi-SSFc, and His6-SSFc. In Figure A, lanes 1 and 2 on the left show the SDS-PAGE results of the supernatant samples of full-length SSF and SSB cell-free protein expression respectively. NC is the control sample with the expression vector. The right picture shows the SDS-PAGE result of the purified proteins. Similarly, Figure B shows the expression and purification results of Ubi-SSFc; Figure C shows the expression and purification results of His6-SSFc. Figure D is a schematic diagram for all proteins involved in this experiment.

### 2.2 ssDNA interaction assay with full-length SSB and SSF

Since the N-terminal of SSF is highly similar to SSB, and SSB is known to have strong interaction with ssDNA, we explored the binding of SSF and ssDNA through gel migration experiments. The 4 nM phage M13 ssDNA plasmid was mixed with 0 nM, 350 nM, and 700 nM protein respectively, and the effects of SSB and SSF on the migration of ssDNA were observed by agarose gel electrophoresis. Experiments showed that both SSB and SSF could bind to different concentrations of ssDNA at a fixed concentration, and the migration rate of the DNA was similar (**Figure 3**). This experiment demonstrates that SSF has similar ssDNA binding ability to SSB.

**Figure 3:**
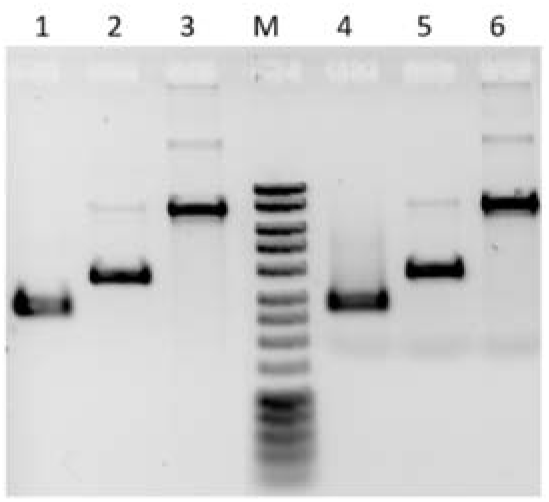
ssDNA binding assay for full-length SSB and SSF. Mix 4 nM of M13 ssDNA template with 0 nM, 350 nM, and 700 nM of SSB (lanes 1-3); repeat the above experiment using SSF (lanes 4-6). Samples were analyzed by 1% agarose gel electrophoresis.

### 2.3 dsDNA binding assay with SSF, Ubi-SSFc和His6-SSFc

In an earlier experiment conducted in our laboratory, we accidentally discovered that the C-terminus of SSF may have dsDNA binding ability. Therefore, we plan to use three different forms of dsDNA, including commercial DNA markers (double stranded structure), purified plasmids, and hairpin structure DNA, to verify the dsDNA binding ability of SSFc (**Figures 4** and **5**). Mix 1 µL of 40 µM SSF or SSF preincubated with 1 µL 40 µM dT35 ssDNA with 1 µL double strand DNA marker, and then load it into 1% agarose gel for electrophoresis detection. Repeat the above experiment using SSB and SSFΔC62. The results show that full length SSF can significantly slow down the gel migration rate of dsDNA marker and can bind ssDNA and dsDNA at the same time, while SSFΔC62 (missing the entire C-terminal sequence of the SSF, only contain the SSF N-terminal OB region) and full length SSB have no effect on the gel migration rate of DNA marker (the left picture in **Figure 4**), indicating that SSF interacts with dsDNA and the interaction region may be located at the C-terminal (SSFc). However, SSB do not interact with dsDNA (consistent with previous reports).The right picture of **Figure 4** shows that the two fusion proteins Ubi-SSFc and His6-SSFc containing only the SSFc region can slow down the gel migration of the double stranded plasmid, and there is no report that ubiquitin and His6 tags interact with DNA. Although the above experiments verified that SSF and dsDNA have a strong interaction, which can significantly slow down the migration of dsDNA in the 3-hour gel electrophoresis, there has not been any report on the accurate binding strength of single strand DNA binding protein and dsDNA. Therefore, we used MST to investigate the dynamic interaction between SSF or Ubi-SSFc and a artificially designed hairpin dsDNA (**Figure 5**). The full-length SSF and Ubi-SSFc both interact with the dsDNA, and the measured dissociation constants (kD) are 45.3 µM and 98.3 µM respectively. As a control, the MST interaction experiment was repeated using full-length SSB, and the results showed that SSB did not interact with the hairpin DNA. The above experiment revealed a never-been-reported feature of a single stranded DNA binding protein, and identified the binding region is the C-terminal non structural region.

**Figure 4:**
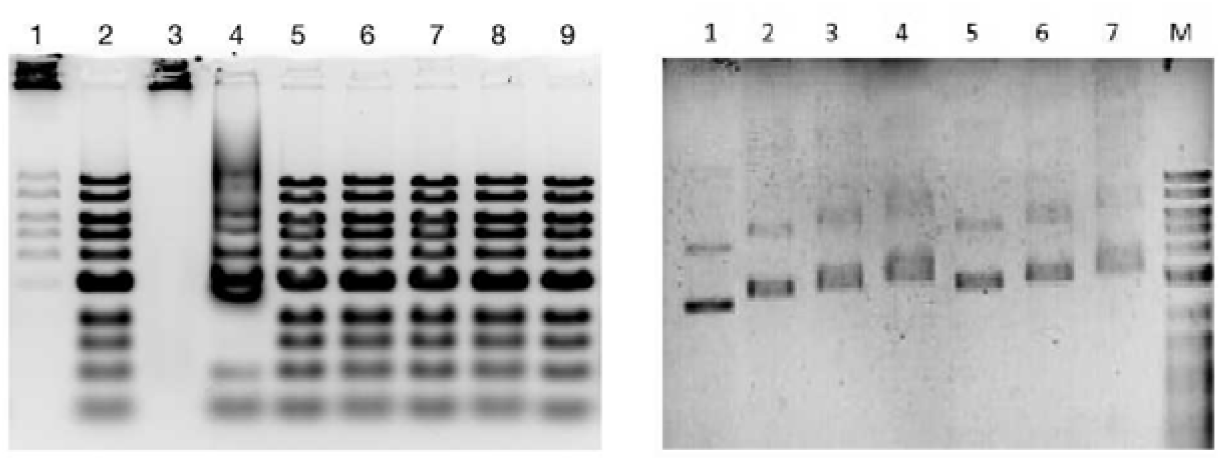
dsDNA binding experiments of full-length SSF, Ubi-SSFc and His6 SSFc. In the left picture lane 1 is the sample mixed with full-length SSF and DNA marker, while lane 2 is the same sample with 0.5% SDS added; lane 3 and 4 contain similar samples as lanes 1 and 2, but with the addition of dT35; lane 5 is the sample mixed with full-length SSB and DNA marker, while lane 6 is the same sample with 0.5% SDS added; lane 7 is the sample mixed with SSFΔC62 and DNA marker, while lane 8 is the same sample with 0.5% SDS added; lane 9 is an untreated DNA marker. In the right picutre, the mixture of 10 ng/µL pET plasmid and 7 µM, 14 µM, and 28 µM Ubi-SSFc were analyzed in lanes 2-4; results of repeating the experiment using His6-SSFc were shown in lanes 5-7; lane 1 is an untreated pET plasmid.

**Figure 5:**
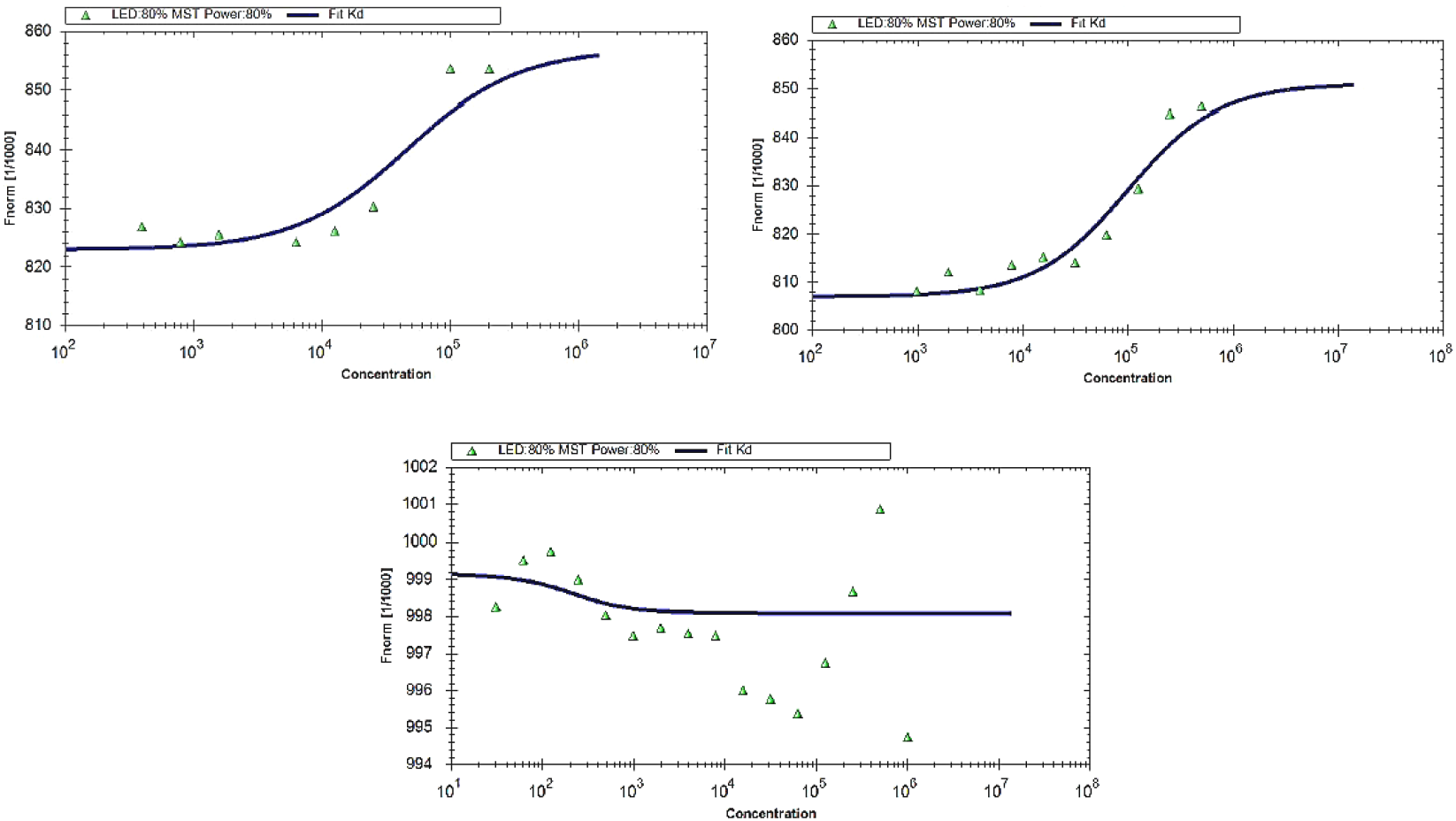
dsDNA binding experiment for full-length SSF and His6-SSFc based on MST. The interaction results between full-length SSF and His6-SSFc with hairpin DNA are shown on the left and right sides of the above picture. The picture below shows the interaction results between full-length SSB and hairpin DNA.

### 2.4 The effect of SSF on intracellular dsDNA

Through gel migration and MST experiments, the dsDNA binding ability of SSF has been confirmed, but its biological significance is still unknown. We extracted the plasmid containing overexpressed SSF and observed SSF’s effect by agarose gel electrophoresis. It was shown that the plasmid disappeared when SSF was overexpressed (**Figure 6**, right). The above experiment was repeated for SSB and His6-SSFc, however no plasmid disappearance was observed. The mechanism by which overexpression of SSF leads to plasmid disappearance may be complex, and we has not conducted further research on this mechanism.

**Figure 6:**
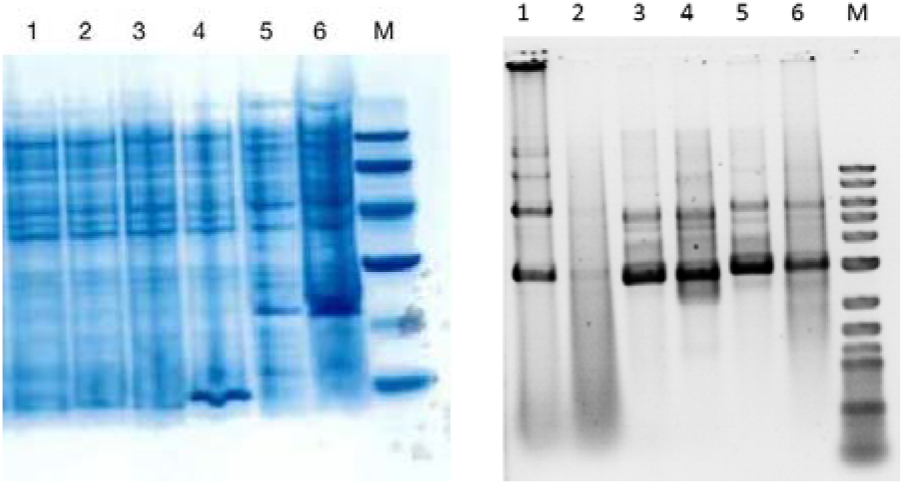
The effect of full-length SSF on double stranded plasmids in *E.coli*. The left picture shows the migration results of proteins in SDS-PAGE. Lanes 1-3 contain before-induction samples of His6-SSFc, full-length SSF, and full-length SSB, while lanes 4-6 contain after-induction samples of corresponding proteins. The right picture shows the migration results of plasmid extracted from *E.coli* with overexpressed proteins in agarose gel electrophoresis. Lanes 1-6 are plasmids from before-induction full length SSF, after-induction full length SSF, before-induction full length SSB, after-induction full length SSB, before-induction His6-SSFc, after-induction His6-SSFc. M is the nucleic acid marker.

## 3 Discussion

Despite its early discovery the presence of another ssDNA binding protein SSF in *E.coli* has not yet been further explored in terms of its structure and functions^[21,22]^. Some early data suggests that SSF is not carried along with ssDNA to recipient bacteria during F-mediated bacterial genetic material exchange^[21]^, and there have been no reports of SSF expression in cells containing F plasmids. It is speculated that SSF is expressed in host cells after bacterial conjugation. But its specific function in recipient cells is still unknown. Although SSF protein is not carried to recipient cells during the binding process, the presence of SSF can reverse and rescue *E.coli* containing temperature sensitive *ssb*-1 mutations^[23]^. Later reports suggested that the ssf gene could also rescue *E.coli* from complete deletion of the *ssb* gene, but *ssf* dependent bacteria may exhibit some abnormalities in DNA replication^[22]^.

By comparing the full-length amino acid sequences of SSB and SSF, we found no similarity in the C-terminus sequences except for the highly conserved OB region at the N-terminus and the eight amino acids at the end of the C-terminus (**Figure 1**). Gel migration and MST test revealed for the first time that the C-terminal sequence had dsDNA binding ability. It is worth noting that the full length SSF can make the dsDNA completely unable to enter the agarose gel electrophoresis, indicating that SSF can combine multiple dsDNA together to form a DNA polymer with large molecular weight (**Figure 4**). However, the SSFc fusion protein that removes the N-terminal OB region does not exhibit strong interaction and multi dsDNA integration ability with the dsDNA marker like full-length SSF. Another interesting phenomenon is that when SSF binds to both ssDNA and dsDNA, there is a clear band blurring in the dsDNA marker after the addition of detergent SDS, indicating that SSF seems to cause recombination or hybridization of certain DNA (**Figure 4**, left lane 3, 4).

This paper also presents the experimental results of SSF causing the disappearance of dsDNA plasmids *in vivo* (**Figure 6**, right figure). So far, reports on SSF have been very limited, and there is a lack of research on the role of SSF in DNA transfer during bacterial conjugation. This paper confirms the binding ability of SSF’s N-terminus to ssDNA and C-terminus to dsDNA, revealing for the first time a single stranded DNA binding protein with dsDNA binding ability. So far, there have been no reports on the dsDNA binding mechanisms of other single stranded DNA binding proteins. It is speculated that the special attribute of SSF is related to the unwinding of dsDNA plasmids, transmission of ssDNA or recombination during bacterial conjugation. More research including gene sequencing and structural analysis are needed to study the unknown activity of SSF in the future to elucidate the function of SSF.

